# MOViDA: Multi-Omics Visible Drug Activity Prediction with a Biologically Informed Neural Network Model

**DOI:** 10.1101/2023.04.07.535998

**Authors:** Luigi Ferraro, Giovanni Scala, Luigi Cerulo, Emanuele Carosati, Michele Ceccarelli

## Abstract

Drug discovery is a challenging task, characterized by a protracted period of time between initial development and market release, with a high rate of attrition at each stage. Computational virtual screening, powered by machine learning algorithms, has emerged as a promising approach for predicting therapeutic efficacy. However, the complex relationships between features learned by these algorithms can be challenging to decipher. We have devised a neural network model for the prediction of drug sensitivity, which employs a biologically-informed visible neural network (VNN), enabling a greater level of interpretability. The trained model can be scrutinized to investigate the biological pathways that play a fundamental role in prediction, as well as the chemical properties of drugs that influence sensitivity. The model leverages multi-omics data obtained from diverse tumor tissue sources and molecular descriptors that encode drug properties. We have extended the model to predict drug synergy, resulting in favorable outcomes while retaining interpretability. Given the often imbalanced nature of publicly available drug screening datasets, our model demonstrates superior performance compared to state-of-the-art visible machine learning algorithms.

## 1 Introduction

Large-scale genomics studies have been instrumental in understanding the recurrent somatic genetic alterations within a cancer cell, including chromosome translocations, single base substitutions, and copy-number alterations [7], and for the characterization of their functional effects in transformed cells [45]. One of the main challenging questions in this field is how to exploit all this molecular information to identify therapeutic targets and develop personalized therapies [17, 49]. Understanding the molecular features influencing sensitivity to drugs is the key element for developing personalized therapies to predict that patients should be treated with specific drugs.[32]. Machine learning models can exploit multi-modal screening datasets to develop predictive algorithms useful to associate omics features with response such as [15] Genomics of Drug Sensitivity in Cancer (GDSC) [51], Cancer Cell Line Encyclopedia (CCLE) [9], Cancer Therapeutics Response Portal [10], NCI-60 [46]. There have been several attempts at utilizing the data from these screenings to various machine learning frameworks such as Variational Autoencoders [42], Deep Networks [13, 41], Convolutional Neural Networks [14, 31], ensemble Neural Network models [48] and a combination of these approaches with different encodings of the features [36] in order to predict the half-maximal inhibitory concentration (*IC*_50_)[8, 13]. Most of these studies use the machine learning models as “black boxes” optimized for prediction accuracy without the possibility of interpreting the biological mechanisms underlying predicted outcomes. However, there is a common need for humans to understand the rules behind model predictions, especially when the final goal is to prioritize drugs (or drug combinations) to be used in clinical trials. Recently, Ideker and colleagues proposed a “visible neural network” to address this issue [27]. The model, called *DrugCell*, encodes cells genotypes into a trainable network constituted by modules organized according to the Biological Process Gene Ontology (GO) hierarchy [5], where each module is associated with a specific GO term and is connected to the nodes (genes) annotated with that specific term. Interpreting the activity of each module allows the association between specific biological pathways and drug response to be discovered.

DrugCell represents one of the first attempts of using *interpretable machine learning* [38] for drug sensitivity prediction. Besides its great novelty, there are several possibilities to extend and improve this biologically informed approach. First, DrugCell relies on somatic single nucleotide variations profiles of the screened models. Second, it is important to consider the imbalanced nature of the data since, in almost all available large-scale screening repositories, sensitivity values tend to be skewed toward values representing a lack of sensitivity, with a small minority representing the sensitivity of a cell line to specific drugs. Finally, it should be important for an explanation tool to allow the subsetting of specific cell lines on which queries can be performed. The use of multiple therapies in patient care has become increasingly popular in recent years, as it has the potential to offer both greater efficacy and fewer side effects [25]. By combining multiple drugs, it is possible to achieve a synergistic effect, in which the combined effect of the drugs is greater than the sum of their individual effects. This strategy can be useful for treating cancer, as it allows for targeting of multiple pathways in the cancer process, resulting in a more comprehensive approach to treatment. This can therefore reduce the doses required for each drug and minimize the potential side effects [2].

The field of drug research has the potential to greatly impact human health. In order to advance the research in this area, we propose a novel approach to drug activity prediction using a Multi-Omics Visible Drug Activity prediction model, or MOViDA. MOViDA extends the existing DrugCell’s visible network approach by incorporating pathway activity from gene expression and copy number variation data. This allows for a more comprehensive understanding of the interactions between drugs and biological systems, leading to more accurate predictions of drug activity. To ensure that the data used to train MOViDA is representative and balanced, the training algorithm employs a random sampler based on a multinomial distribution. This helps to account for skewness in the input dataset. In addition, MOViDA also enhances the interpretability of drug descriptions by using fingerprints and molecular descriptors. These descriptors relate the 3D molecular structure of drugs to their physical-chemical and pharmacokinetic properties, making it easier to understand the impact that a drug may have on a biological system. The results of our study show that MOViDA outperforms existing models in predicting drug sensitivity, particularly for positive treatments. To further enhance the biological interpretability of the model, we developed an *ad hoc* network explanation method to score the pathways affecting sensitivity prediction in specific sets of cell lines. Finally, we have also extended MOViDA to make drug synergy predictions.

## 2 Methods

### 2.1 Network Architecture

MOViDA is a feedforward deep neural network that predicts the drug sensitivity of a cell line. Different omics assays represent each cell line. The structure of the whole Neural Network is separated into two branches (Figure 1a): a Visible Neural Network (*VNN*) and a feedforward Artificial Neural Network *ANN*. The *ANN* on the right branch is a neural network taking as input a combination of PubChem fingerprints [1] and molecular descriptors relating 3D molecular shape with physical-chemical and pharmacokinetic properties [16]. Drug features are encoded into a three-layers neural architecture with 100, 50, and 6 nodes. The *VNN* (Figure 1b) on the left branch represents a Biological Process Gene Ontology hierarchy composed of five layers, one element as a root, and a total of 2086 GOs. Each GO is connected to more generic GO ancestors (at least one) and is represented by a sub-submodule composed of a set of *k* + 1 nonlinear units.

**Figure 1:**
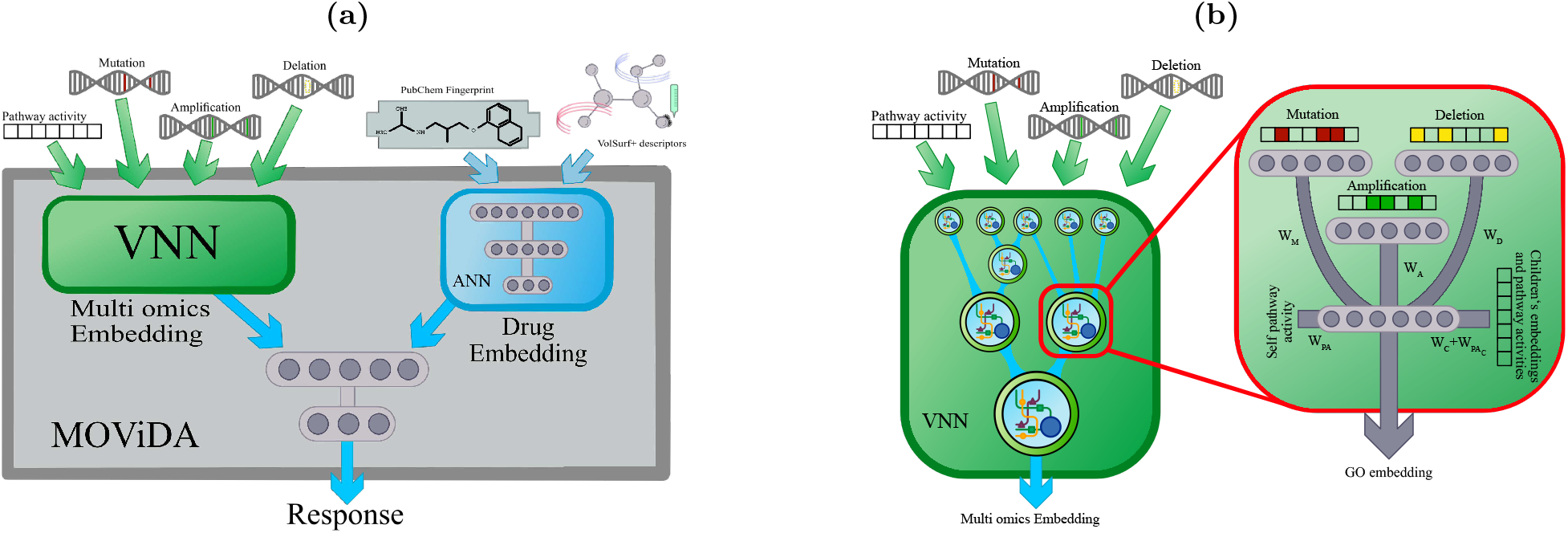
MOViDA architecture. a) The network is composed of three distinct sub-networks. The Multi-omics Embedding net takes in input multi-omics profiles of a cell line model. In contrast, the Drug Embedding net receives the drug description, composed of PubChem fingerprints and VolSurf+ molecular descriptors. The final layers combine the embeddings and predict the AUC. b) The Multi-omics Embedding net comprises a set of modules, each representing a specific Gene Ontology term. The modules are connected according to the Biological Process Gene Ontology Hierarchy. Each GO sub-module takes in input the multiomics profile of a cell line model, considering only the genes associated to former.

*k* units are connected to the input layer and the output of previous layers. Each unit also receives a normalized gene set enrichment score (NES) of that GO term computed from gene expression. This value is concatenated with the activation of the *k* units and fed to the next layer in the hierarchy. The input layer is composed of nodes of three different kinds: mutations, amplification, and deletions. Each GO sub-module is connected to the input genes annotated with that term.

The activation of the units at the root of the hierarchy represents a multi-omic embedding of the cell line. The training phase aims at learning the weights of each subsystem. In particular, every unit of each module *s* has the following output:

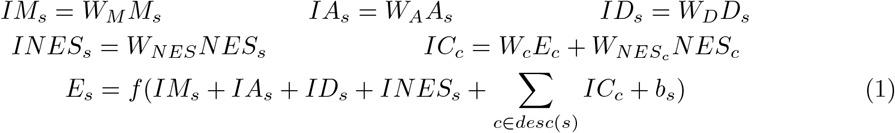

where: *M*_*s*_, *A*_*s*_, and *D*_*s*_ are the binary vectors that describe the mutation, amplification, and deletion status of the genes associated with the subsystem *s* and *W*_*M*_, *W*_*A*_ and *W*_*D*_ are the corresponding weights; *W*_*NES*_ is the weight of the normalized enrichment score *NES*_*s*_ of the term *s* resulting from gene expression; *W*_*c*_ and 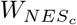 are the d embedding *E*_*c*_ and *NES*_*c*_ of child *c* of the considered subsystem. *E*_*s*_ is the embedding of a subsystem *s*, which is a nonlinear transformation *f* of the inputs consisting of hyperbolic tangent and batch normalization, and *b*_*s*_ is the bias term.

A third neural network combines the multi-omics embedding with the drug features embedding and predicts the cell’s response to the drug, measured as the area under the dose-response curve (AUC). During the training phase, the input data was split into three sets: training (80%), testing (10%), and validation (10%) sets.

### 2.2 Datasets

We used the Genomics of Drug Sensitivity in Cancer database (GDSC) [51] and the Cancer Therapeutics Response Portal v2 (CTRP) [10] to collect 383,998 triplets representing cell line, drug, and cell survival after treatment measure as AUC value. Overall, our dataset contains 889 cell lines and 684 drugs. Each drug is represented by 1009 variables, 881 molecular fragments from PubChem fingerprints [1], and 128 molecular descriptors from the software VolSurf+ [16], as detailed in Table S1. To represent the molecular properties of a cell line, we use the mutation and copy number profiles stored in three binary vectors, where the value corresponds to the presence or absence of a mutation/deletion/amplification in a particular gene in a given cell line, which were downloaded from the GDSC data portal [51]. We selected 4870 (top 2.5%) frequently mutated genes in cancer using the pan-cancer compendium encompassing 33 cancer types and more than 10,000 tumor-normal exome pairs [19]. Analogously, 2612 and 3625 genes contained in focal recurrently amplified copy number segments and deleted copy number segments respectively, selected as described in [24]. These genes were further filtered for those associated with at least one GO term present in the MOViDA hierarchy, obtaining 2931 and 2097 genes for amplifications and deletions, respectively. Gene expression was also used to compute a normalized enrichment score (NES) using single-sample gene set test using the Mann–Whitney–Wilcoxon Gene Set test (mww-GST) available in the yaGST package [22]. NES is an estimate of the probability that the expression of a gene in the geneset is greater than the expression of a gene outside this set: 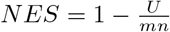 where *m* is the number of genes in a gene set, *n* is the number of those outside the gene set, *U* = *mn* + *m*(*m* + 1) − *T*, and *T* is the sum of the ranks of the genes in the gene set.

For drug combination, we used the Therapeutic Target Database (TTD) [52] to identify potential synergies among drug targets and then used the dataset of pharmaceutical synergies specific to breast, colon, and pancreatic cancer cells created by Jaaks et al. [25] for validation. This approach allowed us to confirm the potential of certain drug combinations to have a synergistic effect on cancer cells and to develop more effective treatment plans, potentially leading to improved outcomes.

To further advance our model’s capabilities, we extended its application to predict drug combination therapies utilizing the dataset presented by O’Neil et al. [40]. We selected the cell lines and drugs with available features, resulting in a dataset of 32 compounds and 32 cell lines, totaling 13376 instances of combined cell line and drug treatments, with 1296 instances considered synergistic. To assess the synergistic interaction between drugs, we employed the Loewe Additivity score [34], utilizing a threshold of 30 to differentiate synergistic from nonsynergistic outcomes. To overcome the limitations posed by the limited size of our dataset, we took steps to reduce the number of input features, specifically by excluding copy number information from the input.

### 2.3 Data Imbalance Strategies

Drug sensitivity data exhibits a significant skewness, characterized by many screens with low sensitivity outcomes (AUC close to 1) and very few with high sensitivity (AUC close to 0). To mitigate the potentially deleterious effects of this data imbalancing during the training, we used a weighted random sampler based on a multinomial distribution estimated from the data.

The AUC sensitivity scores are divided into twelve equally spaced bins between 0 and 1.2, and we used the inverse frequencies with additive smoothing to fix the weights of the multinomial sampler:

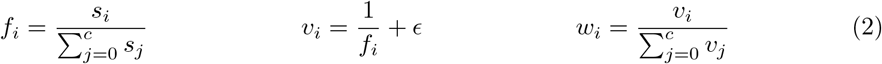

Where *s*_*i*_ is the number of samples in bin *i, c* is the number of bin, *f*_*i*_ is the relative frequency of the bin *i, ϵ* is the smoothing penalty term.

We also used a *weighted loss* function to give more importance to errors associated with lower scores of ground truth and predictions. Hence we adopted the following *double-weighted MSE loss*.

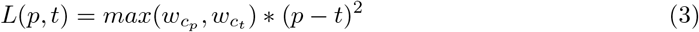

Here, *p* is the prediction of a model, *t* is the ground truth, *c*_*p*_ and *c*_*t*_ are the corresponding bins and 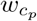 and 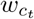 are the weights associated with these bins as computed in Equation (2). This loss function guarantees higher weights for errors when either the ground truth or the prediction are in a class with few samples and, at the same time, lower weights for predictions when they are far from the ground truth.

If an evaluation measure conceived for balanced datasets was used, a trivial system assigning the majority of the items the values with the highest frequency may even outperform genuinely engineered systems. Evaluating measures that can handle imbalance are recommended to avoid these cases. We divided the values of the target function (the drug sensitivity) into 12 discrete bins in the interval (0,1.2) and considered each bin as a separate class, and adopted a measure developed in the field of ordinal regression [6]. This is motivated by the fact that sensitivity classes can be considered ordinal variables, and the ordering between the values is significant, as they represent discretized degrees of sensitivity. A simple and efficient approach to measure the performance in ordinal regression tasks for imbalanced datasets is the *macroaverage MSE* which is based on a sum of the classification errors across classes.

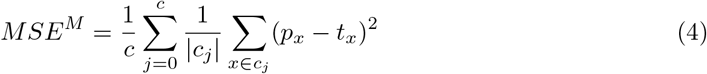

where *c*_*j*_ represents the set of samples in class *j, c* is the number of classes, *t*_*x*_ id the ground truth of sample *x* and *p*_*x*_ is its prediction. The macroaverage MSE does not depend on the frequency of each class, as every class contributes to 1*/c* of the total measure. Therefore trivial assignments are penalized, whereas to have better *MSE*^*M*^ the errors in all classes should be minimized.

### 2.4 Model Explanation

The Biology informed nature of MOViDA, as well as of DrugCell [27], P-NET [20], and PASNet, [23] allows performing accurate post-hoc analyses exploiting the above methods. This enables us to identify the biological processes that contribute to the prediction of a cell line’s drug sensitivity the most. The state-of-the-art methodologies, such as LIME [43], DeepLIFT [47], DeepExplain [4], SHAP [35] implemented in the *Captum* library [26], are not completely suited to our case since most of them allow us to measure the contribution of either an individual input node or a full layer. Instead, our visible network is composed of sub-modules. Therefore, we developed an interpretation score, relative improvement score (*RIS*), specifically tailored for our model that measures the relative contribution of a sub-module concerning its children in the GO hierarchy. This score is inspired by *ablative brain surgery* [37], which involves removing certain components of the brain while keeping all of its functions intact.

First, we calculate the prediction for a specific drug-cell line pair. Then we recalculate the prediction after silencing the output of each sub-module one by one, setting weights and biases to 0. Similarly, we silence all children subsystems for each GO and obtain the third prediction. In the case of leaf nodes, we silence the corresponding inputs. The *RIS* score expresses the importance of a term during the prediction phase and its ability to combine the information from its children, comparing the deviations from the actual prediction of the models ablating first the father and then its children terms. The *RIS* is computed as follows:

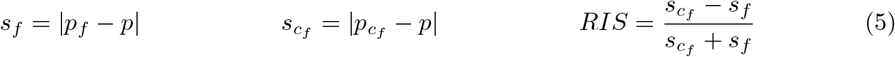

where *p* is the effective prediction for a specific drug-cell line pair, *p*_*f*_ and 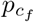 is the prediction obtained by silencing a GO subsystem and its children, respectively. *RIS* is the interpretation score. For a given GO subsystem, positive RIS values correspond to a larger deviation in predictions when silencing children compared to the father.

The advantages of *RIS* over the score adopted in DrugCell (RLIPP) are that: i) it can be calculated for each individual drug-cell line pair; ii) there are multiple ways to aggregate these values, by drug or by specific cell line types.

To further investigate the model, we have inspected all the elements that compose the inputs describing the drugs. The importance score of each feature was performed using DeepLift, whose implementation is based on the algorithm of Shrikumar et al. [47] and gradient formulation proposed by Ancona et al. [4].

### 2.5 Drug Combination Strategies through Relevant Subsystems

The synergistic effects of multiple drugs can have a profound impact on pharmacological interventions for cancer [3]. By combining multiple drugs, it is possible to harness cancer’s resistance to particular anticancer drugs, and create a more effective treatment plan. A recent effort toward understanding the combined effects of drugs has been carried out by Jaaks et al. [25] by creating a huge dataset of pharmaceutical synergies specific to breast, colon, and pancreatic cancer cells and demonstrating how the context influences the outcome of therapy.

Taking into account a drug and a cell line evaluated in the combination dataset, we used the RIS score calculated from a drug/cell-line pair. We selected the top 5 enriched GO terms along with associated genes. From the collection of drug targets *Therapeutic Target Database* [52], we determine the drugs targeting the genes associated with the previous selection of GO terms, marking them as potentially synergistic for that drug-cell line pair.

We then compared our predictions on the synergistic dataset, marking the right (TP) and wrong (FP) combinations and comparing the ratio of TP to FP and the ratio of synergistic drugs to non-synergistic drugs to understand if the former was significantly higher than the latter. We took all drug combinations studied for a specific cell line and drug and counted how many of these combinations were synergistic (S) and how many are not (NS). We applied the binomial test on TP over (TP + FP), which is the Precision metric, with probability equal to S/(NS+S), thus accounting for the number of synergistic combinations. The p-values for the binomial test and the enrichment scores 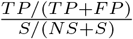 of the above-described tests are used in the volcano plots reported in the Results section.

### 2.6 Extension for Drug Synergy Prediction

We have extended the architecture of our model to see its actual usefulness in predicting synergistic effects of drug combinations as well. For this purpose, we exploited the concept of Siamese neural networks, which involves the use of identical subnetworks, in this case the right branch of our model, that embed drug features. The final ANN concatenates the cell line and drugs embedding. In this way, the order of past drugs is important, so we doubled the initial dataset by considering the two possible combinations of drug pairs. In addition, the problem was set up as a binary class prediction, distinguishing the cases where there is synergy or not. Given the presence of imbalance in the dataset, we used focal loss [30] with hyperparameters *α* and *γ* set to 0.4 and 2, respectively. Additionally, we employed the weighted random sampler as previously described, with a weight-balancing factor of *ϵ* = 5.

## 3 Results

### 3.1 Dataset Imbalance

MOViDA is trained to predict the response of a cellular model (represented by its molecular profile) to a specific drug, measured as area under the curve (AUC). AUC combines information about the potency and efficacy of the drug into a single measure [21]. A value close to 0 means high sensitivity, a value close to 1 represents no effect of the drug, if higher than 1, the drug has the effect of promoting cell viability. Besides the high interest in accurate predictions for drugs with high sensitivity, the majority of drug screens present AUC values that are close to 1, therefore, presenting a marked skewness. Figure S1a shows the distribution of the AUC values present in the dataset after binning AUC values into 12 bins (ten bins for the interval between 0 and 1 and other two for values greater than 1): class 0 (AUC scores in the range [0.0, 0.1]) is 80 times less populated than class 9 (scores in the range [0.9, 1.0]). To mitigate this effect, our approach considers a weighted random sampler and a double-weighted loss (Section 2.3).

Both use the weights calculated as a function of the inverse frequencies of each class plus a smoothing term *ϵ*. Figure S1b shows the number of samples for all classes: besides the raw case (no weights), different scenarios are depicted by varying the *epsilon* parameter that affects the weights. The ideal scenario lies between the raw case and the perfectly balanced dataset (with *epsilon* set to 0), which, on the contrary, could produce too many sample repetitions. After parameter tuning, we chose the value of *ϵ* to 80 as a good compromise, producing, on average, a four-fold repetition for the samples in a less represented class.

### 3.2 Performance and comparison with DrugCell

The accuracy of the prediction was evaluated by measuring the Spearman and Pearson correlation between the predicted AUC values and the actual ones, averaged over 5-fold cross-validation. To assess the issue of imbalance, 100 samples from each class were sampled, and this process was repeated for 1000 runs. The results show that both Pearson and Spearman correlation were 0.89. The DrugCell model showed good results as well, with Pearson correlation of 0.86 and Spearman correlation of 0.88.(Table S2). If we use macroaverage MSE (MMSE), which accounts for the imbalance of the classes, we observed that MOViDA has a lower error (0.025 vs. 0.035) after the cross-validation. Indeed MOViDA can make much more accurate predictions of the AUC in classes with fewer training examples (classes between 0 to 5). Those classes are the most meaningful ones as they represent cases of high sensitivity to drugs. Notably, MOViDA exhibits higher accuracy in these classes compared to DrugCell, with a significantly lower error rate (0.032 vs. 0.060). Conversely, for classes with a larger number of training examples (classes 6 to 11), MOViDA and DrugCell perform similarly, with comparable error rates (0.020 vs. 0.018). We show in Figure 2b the lowest, mean, and highest MSE calculated for each class, averaged across five cross-validation folds, which depicts a specific trend, where MOViDA performs slightly worse for the upper classes (6-11) than DrugCell, but significantly better for the lower classes (0-5). To better explain this behavior, by recasting the regression the sensitivity values as a classification problem in terms of prediction of AUC interval classes, the confusion matrix in the Figures S2a and S2b show that DrugCell tends to over-estimate the majority class. In contrast, MOViDA reaches better accuracy along the cells on the diagonal.

**Figure 2:**
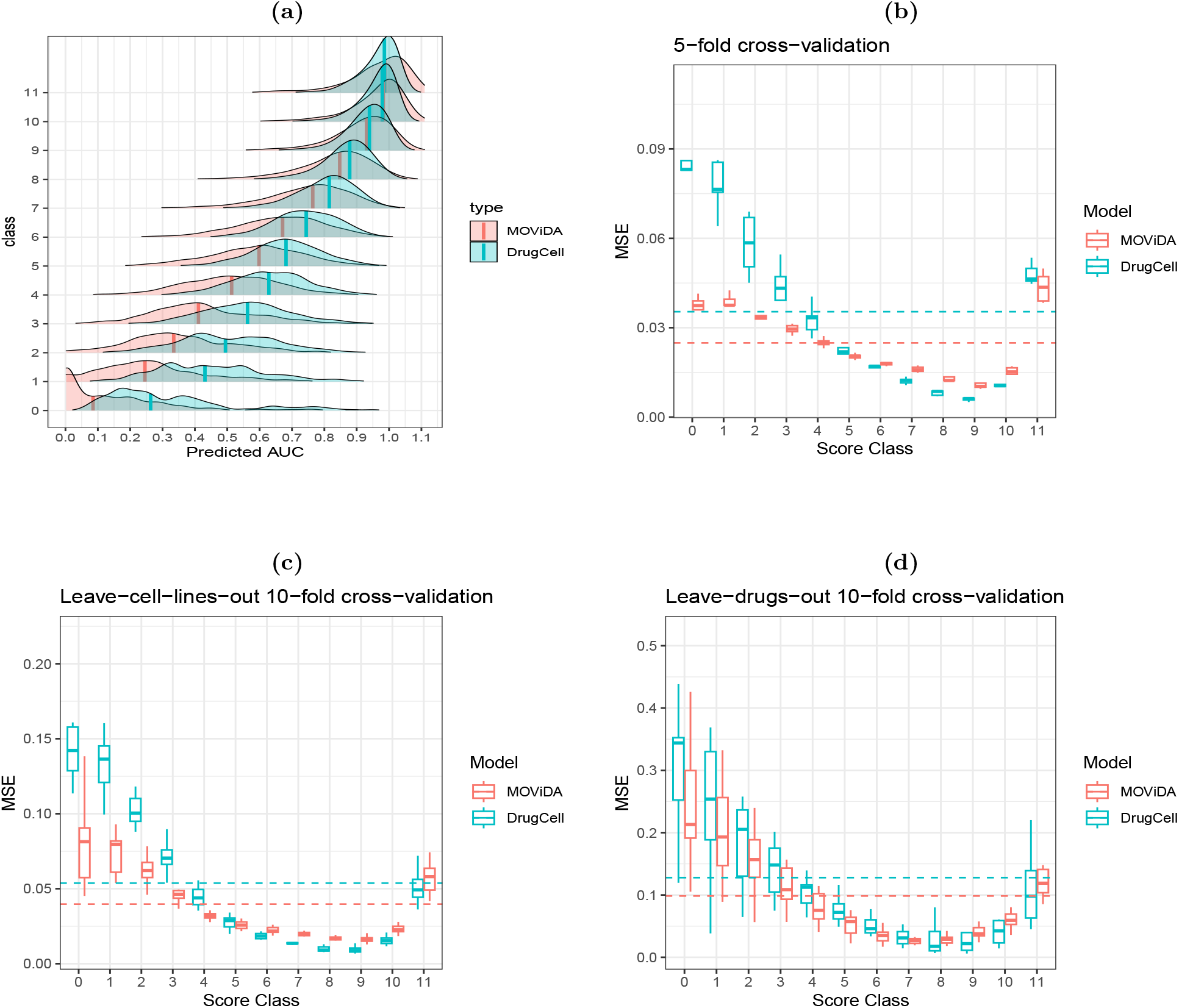
Evaluation and comparison. a) Distribution of predicted AUC compared to the ground truth classes. b) 5-fold cross-validation results for each model’s mean squared error (MSE) per class. The plot illustrates the lowest, mean, and highest MSE values obtained. The dashed lines correspond to the macroaverage MSE averaged over cross-validation. c-d) 10-fold cross-validation in which each fold comprised a unique set of 10% cell lines or drugs that were not included in the other folds.

We investigated the robustness of each model using two cross-validation strategies: leave-celllines-out and leave-drugs-out, both implemented in a 10-fold nested format. We created 10 folds for each strategy, ensuring that each fold contained cell lines or drugs not present in the other nine folds. These cross-validation strategies enabled us to assess the ability of the models to generalize to unseen data and evaluate their accuracies. As reported in Table S2 and Figures 2c and 2d, the results demonstrate that while all models experience a slight drop in accuracy during leave-cell-lines-out cross-validation, MOViDA consistently outperforms DrugCell (0.040 vs. 0.054). In contrast, during leave-drugs-out cross-validation, MOViDA remains stable across the 10 folds, while DrugCell exhibits significant variability, suggesting that it may rely heavily on the drugs in the training set.

The performance of MOViDA was further evaluated (Figures S3) through comparative analyses, varying the type of drug representation used as input (Morgan Fingerprint, PubChem Fingerprint, and VolSurf+ descriptors). Results indicated that MOViDA exhibited the lowest macroaverage mean squared error (MSE) compared to other models, particularly with smaller errors in the lower classes. Furthermore, we have tested MOViDA by replacing the artificial neural network (ANN) for drug embedding with another one that contained more than five times the number of parameters, with 512, 128, 32, 8 nodes respectively, and 4 linear layers. Our results showed that MOViDA, despite having fewer parameters, generalized better, mainly in the lower classes. Finally, MOViDA was compared with a Multi Layer Perceptron (MLP) consisting of 5 linear layers (1024, 256, 64, 4, 1 nodes each) with ReLU activation functions to assess whether the trade-off between explainability and performance exists. The results indicated that MOViDA and the standard network performed similarly, with comparable macroaverage MSE.

### 3.3 The RIS score identifies pathway dependencies in specific cellular models

We have implemented a novel score called relative improvement score (RIS), which is based on ablative brain surgery [37]. Our score is calculated by determining the deviation from the prediction for each node, caused by removing: the node layer in the VNN and the deviation from the prediction; all its children nodes. This helps us assess the contribution of the parent node to the overall prediction compared to its children. This score also has the advantage of being calculated for each specific cell line-drug prediction, so we can show which GOs are most predictive for a specific case or tissue (represented as a group of cells) or drug.

Among the leukemia cell lines, we selected the ALLSIL cell line highly sensitive to GSK1070916, an ATP-competitive inhibitor of Aurora kinase, which is important during cell division. The RIS scores associated with this prediction revealed that *anion transmembrane transport* (GO:0098656) is among the most important modules for prediction (Figure 3a). The overexpression of ATP-binding cassette (ABC) transporters, particularly ABCG2, contributes to reduced cytotoxicity of GSK1070916 [50]. The family of these genes is responsible for transporting substances across the cell membrane using the energy produced by ATP electrolysis. Interestingly, ALLSIL is ABCC9 mutant which, together with ABCG2, is downregulated in this cell line. Similarly, *proteolysis* (GO:0006508) has a high RIS score for this cell line-drug pair. This can be attributed to AURKB (aurora kinase B) phosphorylating caspase-2 by mediating its proteolysis [29]. As a result, cell division is not stopped. In our case, GSK1070916 inhibits AURKB promoting apoptosis of the cancer cell. AURKB is over-expressed in the ALLSIL cell line.

**Figure 3:**
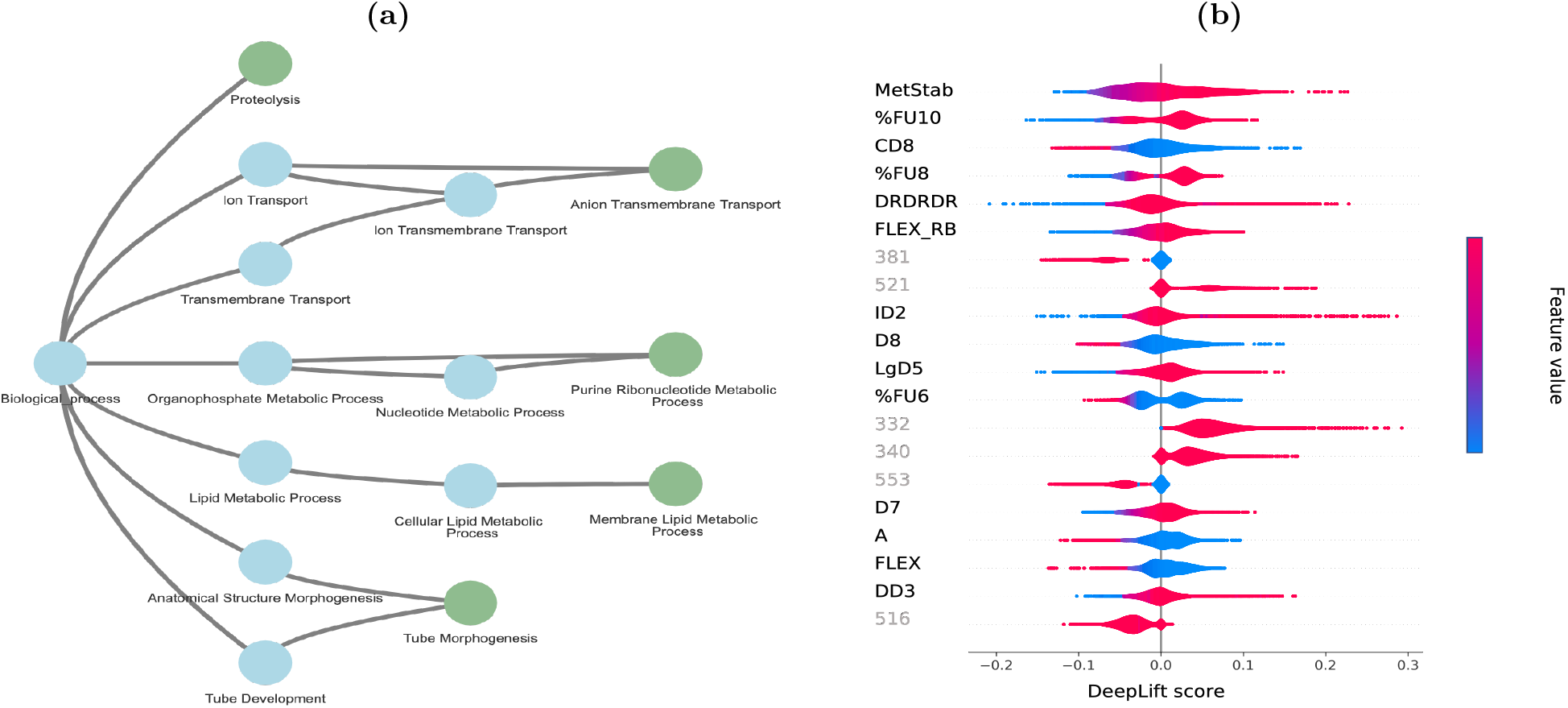
Explainability. a) Top 5 RIS score associated to GOs (green nodes), considering the ALLSIL cell line and GSK1070916 drug. The whole sub-tree (blue nodes) is displayed. b) Deep Lift drug feature interpretation of Liver tissue. The color represents the feature value for a specific prediction. If feature values are low (blue) on the left and high (red) on the right of the violin, AUC is directly dependent on the feature. Feature names are colored differently if they are VolSurf+ descriptors or PubChem fingerprint bits in black and gray, respectively.

An high RIS score is also reported for the *positive regulation of the reactive oxygen species metabolic process (ROS)* pathway (GO:2000379) associated with the DB cell line (lymphoma) when administered with Dinaciclib (an inhibitor of CDK1, CDK2, CDK5 and CDK9). Interest-ingly, it has been recently reported that the inhibition of CKD leads to increased mitochondrial ROS levels, confirming this pathway’s importance in the cellular response to this exposure [44].

### 3.4 Drug features interpretability

As a complementary interpretation step, we can also measure the impact of individual drug features on the model’s predictions utilizing DeepLift score [47]. Figure 3b shows the 20 most important features of our model. The importance lies in the variability of the score for the various cell lines: the more it varies, the more significant it is for predictions. The most relevant feature is the VolSurf+ descriptor METSTAB for all the cell lines. Such descriptor refers to metabolic stability (measured on human liver microsomes), mostly due to isoform 3A4 of the cytochrome P450 system. We have noticed a direct relationship between such a feature with the AUC. This means low values for metabolic stability (thus, fast CYP3A4-mediated metabolism) for high-sensitivity drugs. This agrees with the absorption, distribution, metabolism, and excretion (ADME) profile of many anticancer drugs, most of which are metabolized in the liver by CYP3A4. Several features refer to drug lipophilicity; among these, the VolSurf+ descriptors D8 and CD8 refer to highly lipophilic regions of the molecules, and characteristics of active molecules (low AUC values).

Two features refer to molecular flexibility, namely the VolSurf+ descriptors FLEX and FLEX RB. Given their lift values, we can argue that for most of the predictions, flexibility is inversely related to AUC, whereas the number of rotatable bonds is directly related to AUC. Although it is uncommon to have an opposite behavior for these two features, an attempt to generalization may be that anticancer drugs are generally flexible but with a low number of rotatable bonds (compared to the overall number of bonds). The VolSurf+ descriptors %FU8 and %FU10 can measure the percent of the unionized fraction at a given pH (8 or 10). According to violin colors, the system identified a direct relationship with AUC; in other words, many anticancer drugs have strong or weak acid groups, that is reflected onto the significant presence of ionized species at basic pH.

### 3.5 Drug Combination predictions

MOViDA predictions can be used to uncover potential drug synergies. Given the interpretation score for specific drug-cell line pairs, we select the genes involved in GO terms with the highest scores and prioritize as potential combinations the drugs targeting these genes (Methods 2.5), reported in Supplementary Material.

The volcano plot in Figure 4a shows the cell line-drug pairs for which the candidate molecules are enriched for experimentally validated synergistic drugs. For example, the synergy predictions associated with the breast cancer cell line JIMT1 and the drug MK-2206, a highly selective inhibitor of Akt1/2/3among, has among the top 5 scoring GO categories the GO:0007169 (transmembrane receptor protein tyrosine kinase signaling pathway) and the GO:0007584 (response to nutrient). Our model selects Lapatinib, PD173074, Axitinib, Linsitinib, Sapitinib, and OSI-027 as potential candidates for combination therapy with MK-2206. They are all tyrosine kinase inhibitors involved in tumor cell growth. The association between these drugs and MK-2206 is well documented in the literature, as many tyrosine kinases are part of the PI3 kinase-AKT cascade, affecting mTOR activity [28]. Another relevant combination consists of Navitoclax and Vorinostat associated with the MDAMB231 cell line. The latter is an HDAC inhibitor, which decreases the expression of BCL2 family proteins, as described by [18]. Since Navitoclax is an inhibitor of this anti-apoptotic protein family, it has been shown that its efficacy, combined with Vorinostat, can induce apoptosis in cancer cells [39].

**Figure 4:**
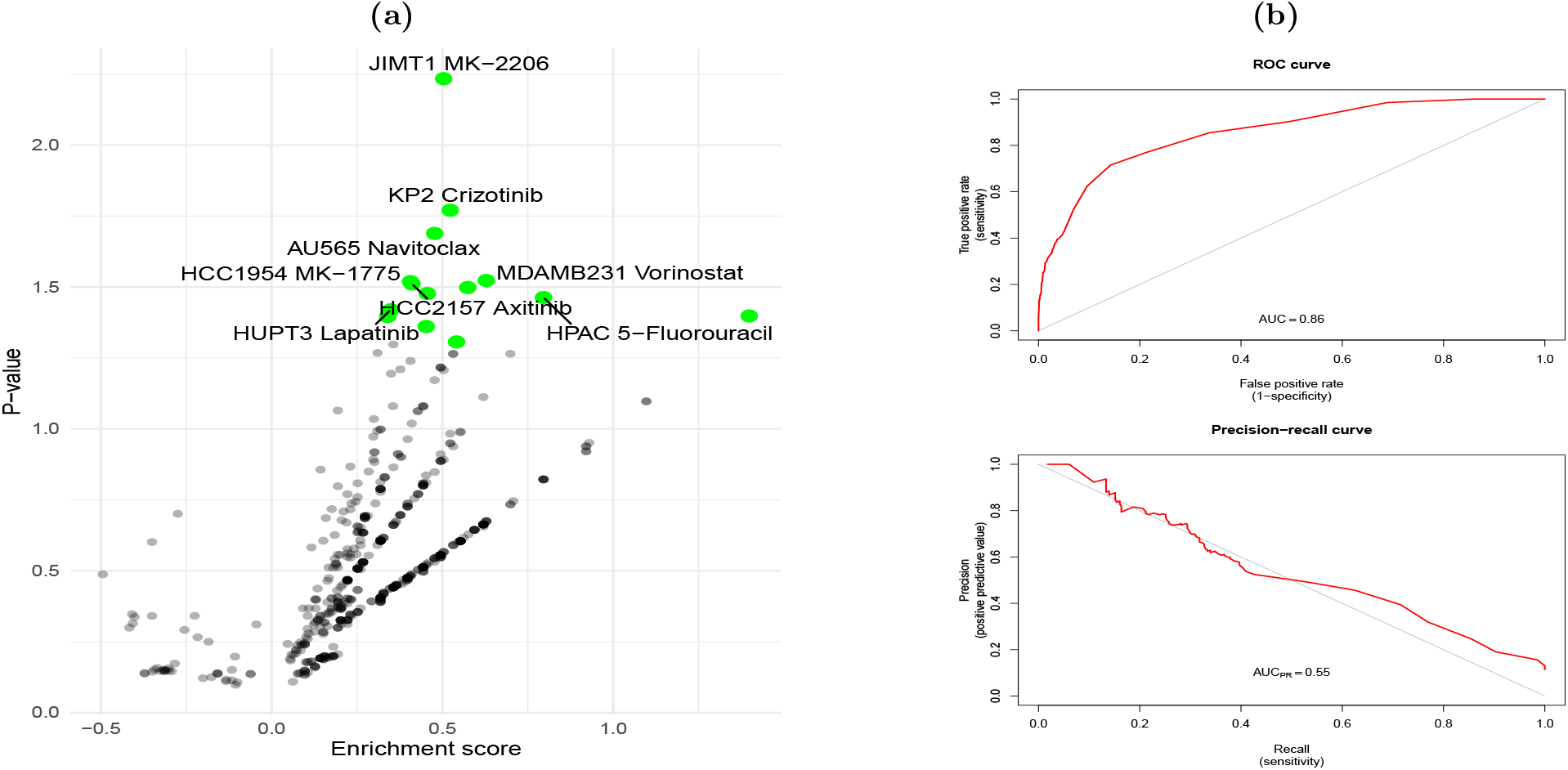
Evaluation of Drug synergy prediction. a) Enrichment scores against the p-values of binomial, testing the Precision of the model, using as the probability the percentage of synergistic combination. The green points correspond to drug-cell lines pairs that have a significant number of positive drug candidates, considering the numerosity of synergies in the dataset. b) AUC scores were calculated for ROC and PR curves, which are graphical evaluations of binary classification model performance. PR shows precision vs recall, while ROC plots TPR (recall) vs FPR. Both help determine the optimal threshold for making binary classification predictions by showing the trade-off.

MOViDA was initially developed as a regressor for predicting drug sensitivity in cancer cells. However, in order to better support the discovery of new and effective cancer therapies, the model has been extended to classify drug synergy as well. To evaluate the performance of the extended model, we used two commonly used metrics in machine learning: area under the Receiver Operating Characteristics (ROC) and Precision-Recall (PR) Curves. The results showed that MOViDA was able to achieve a ROC of 0.86 and PR of 0.55 (Figure 4b), which indicates high accuracy in classifying drug synergy despite the imbalanced nature and small size of our dataset, which is often a challenge in drug discovery research. We conducted a comparative analysis between our model and a standard neural network consisting of 4 linear layers with batch normalization, trained using identical hyperparameters to MOViDA. Surprisingly, our model yielded superior performance, with both areas exceeding those of the compared model (Figure S4).

## 4 Conclusion

In this manuscript, we presented MOViDA, a biologically inspired neural network architecture for the prediction of drug sensitivity of cellular models of cancer. The assessment of anti-cancer drugs and the identification of potential synergistic effects can be ideally assessed by using patient-derived cell lines [33]. However, this process requires substantial time, and there is no guarantee of efficiency. The use of machine learning (ML) to exploit the variety of screening data already available, together with the knowledge of the molecular features of cellular models, can help to accelerate the process of drug prioritization for experimental validation [17] and candidate combination therapies [25]. The adoption of a biologically informed architecture has three main advantages: a) it allows to uncover the role of specific pathways in response to drug stimuli; b) it improves the trust in predictions, especially among non-ML experts and c) the efficient parameterization of our model can simplify the learning process rather than use arbitrarily overparameterized, architectures for prediction, simplifying interpretability. Most drug sensitivity prediction models only use gene expression data [12], however, the effect of single nucleotide mutations, DNA methylation and DNA copy number variation on drug sensitivity should also be considered. Here we have presented a visible neural with an improved accuracy level due to the use of multiple omics platforms and the better handling of imbalance of data. We also have developed an interpretability score that has the advantage of producing a value for every cell line-drug pair and, therefore, can be summarized in terms of cellular models derived from the same tissue/cancer subtype or at the level of individual drugs. We have shown that our score produces meaningful results that can be the subject of experimental follow-up. We have also introduced a set of features that can be directly related to behavior or chemical groups. We confirmed the importance of chemical features such as LogP, FLEX as well as %FU(4-10) also observed in the inhibition of glycoprotein [11]. In conclusion, the model has been successfully extended for drug synergy prediction while maintaining its interpretable nature.

## Appendix A Key points

- The adoption of a biologically informed architecture allows to uncover the role of specific pathways in response to drug stimuli and it improves the trust in predictions.
- Machine learning methods aimed at predicting sensitivity to drugs should take in account the unbalanced nature of large-scale drug screening datasets.
- Molecular drug features and multi-omics embedding can significantly improve the accuracy of sensitivity predictions.

## Appendix B Availability and implementation

MOViDA is implemented in Python using PyTorch library and freely available for download at https://github.com/Luigi-Ferraro/MOViDA.

RIS score and drug features are archived on Zenodo https://doi.org/10.5281/zenodo.7406327

## Appendix C Funding

The research leading to these results has received funding from AIRC under 5 per Mille 2018 ID. 21073 project – P.I. Maio Michele, G.L. Ceccarelli Michele. The research leading to these results has received funding from Italian Ministry of Research Grant PRIN 2017XJ38A4 004 and Associazione Italiana per la Ricerca sul Cancro (AIRC) IG grant 2018 project code 21846 P.I. Ceccarelli Michele.

## Appendix D Competing interests

No competing interest is declared.

## Appendix E Author contributions statement

L.F. and M.C. conceived the study, L.F. developed the code and generated the results, G.S., L.C. and E.C. analysed the results, L.F., E.C. and M.C. wrote the manuscript, all authors reviewed the manuscript.

## Appendix F

## Acknowledgments

We thank Associazione Italiana per la Ricerca sul Cancro for the support. We thank Molecular Discovery Ltd. for supplying the VolSurf+ software (https://www.moldiscovery.com/software/vsplus).

---

**Figure S1:**
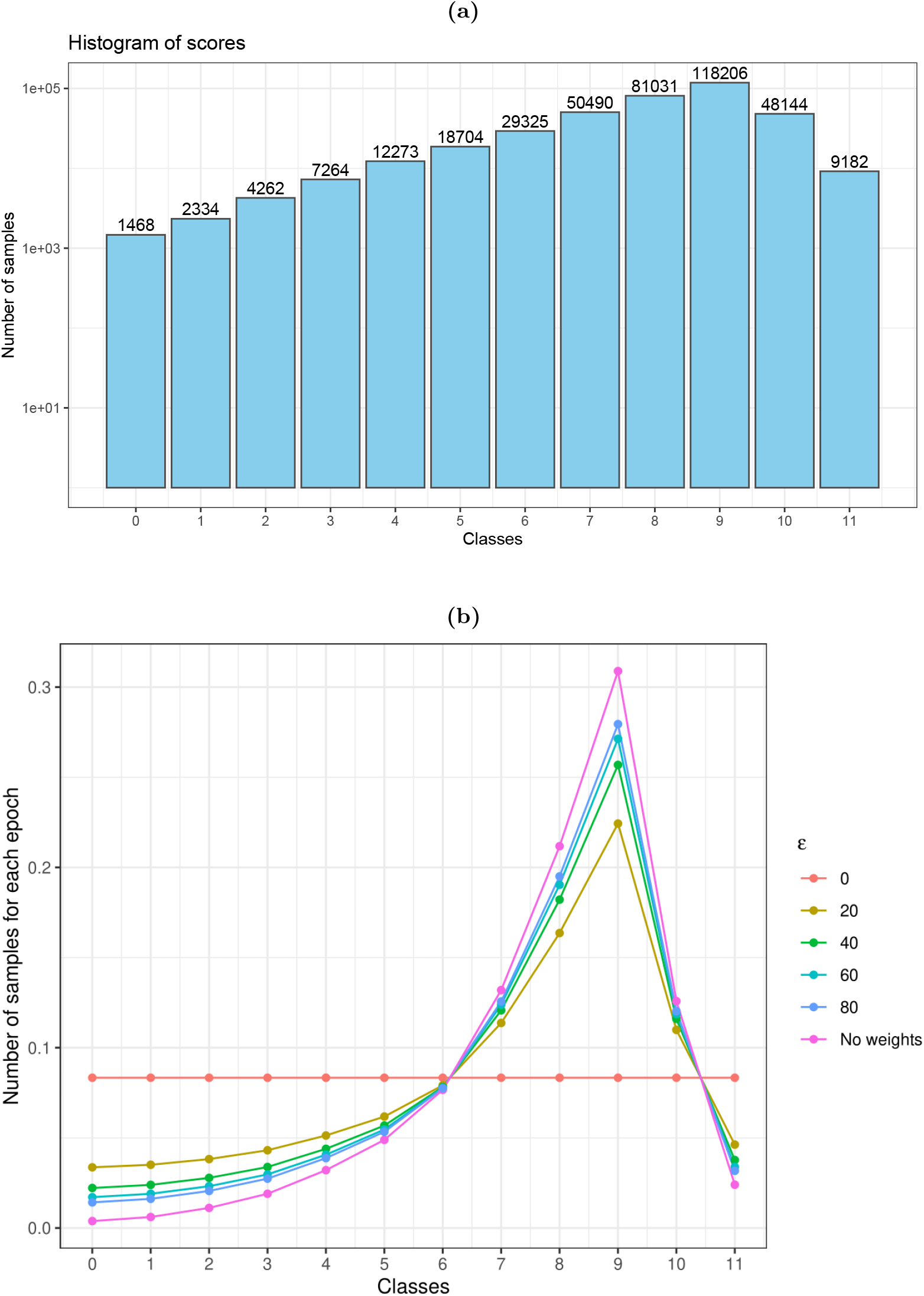
Data Imbalancing review. a) BarPlot distribution of AUCs divided in 12 classes, showing high data skewness, with very few examples for classes corresponding to high drug sensitivity (low AUC values). b) Probability to pick up a sample of a specific class by varying *ϵ* parameter. It is computed as *w*_*i*_ * *s*_*i*_ normalized.

**Figure S2:**
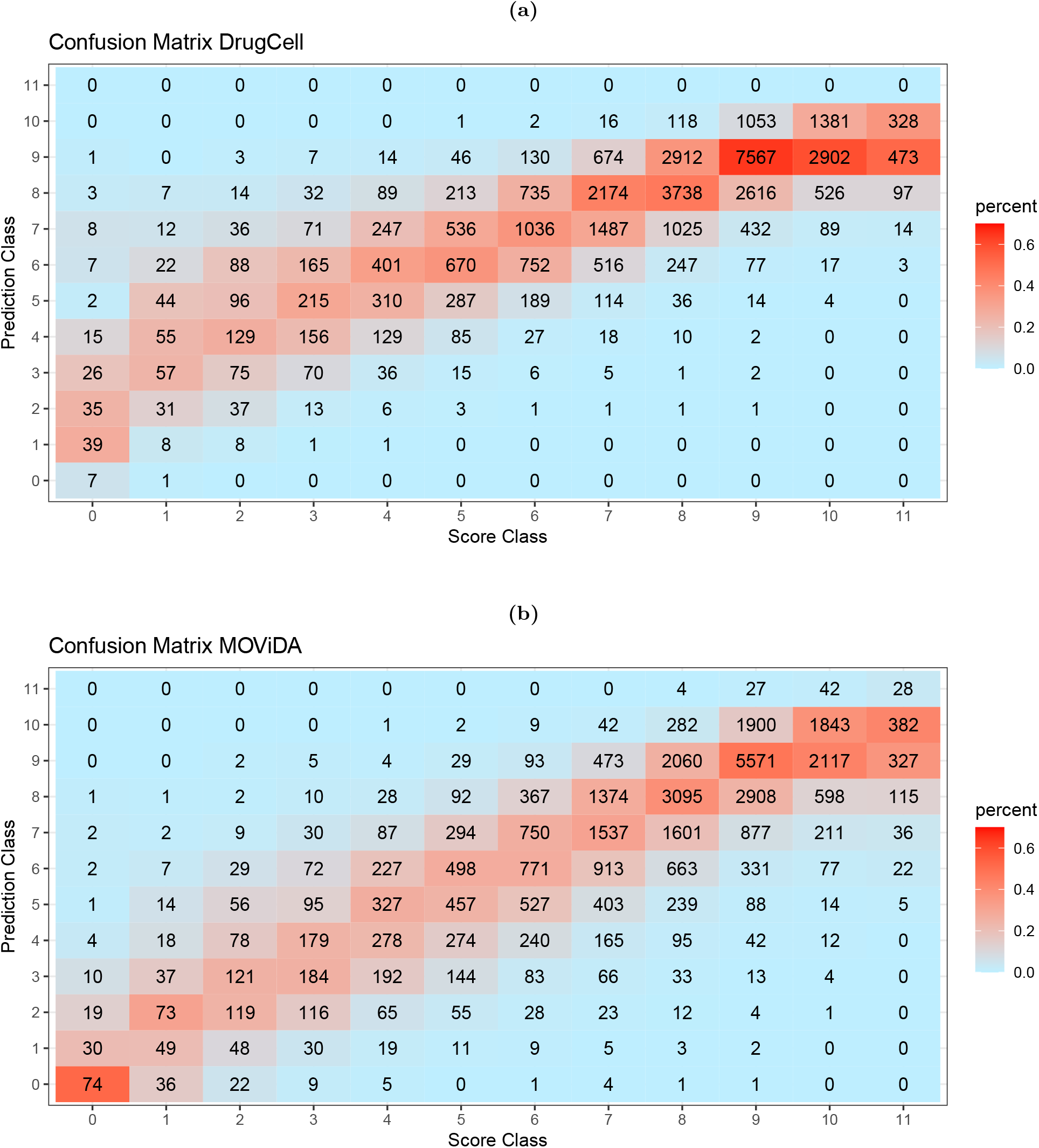
Evaluation and comparison. Confusion matrices of DrugCell and MOViDA respectively, the percentage of each box is calculated based on the numerosity of the reference score class

**Figure S3:**
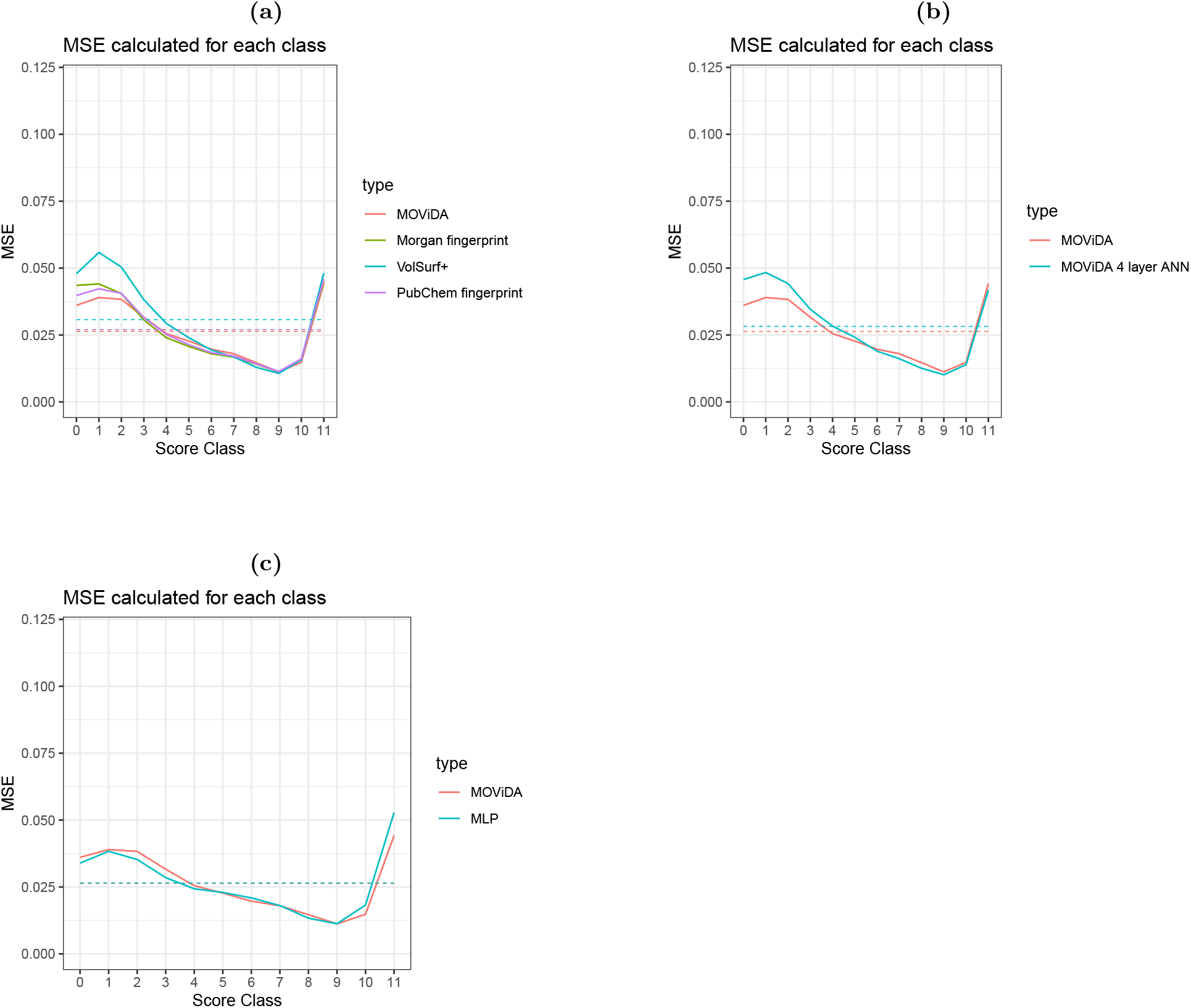
Evaluation and comparison. MSE computed for each class with dashed line correspondind to the macroaverage MSE. Comparison of MOViDA: a) varying drug input (Morgan fingerprint, VolSurf+, PubChem fingerprint; b) substituting the ANN for drug embedding with 4 linear layers ANN of 512, 128, 32, 8 nodes respectively; c) with a MLP, composed of 5 linear layers, with 1024, 256, 64, 4, 1 nodes respectively, and ReLU as activation functions.

**Figure S4:**
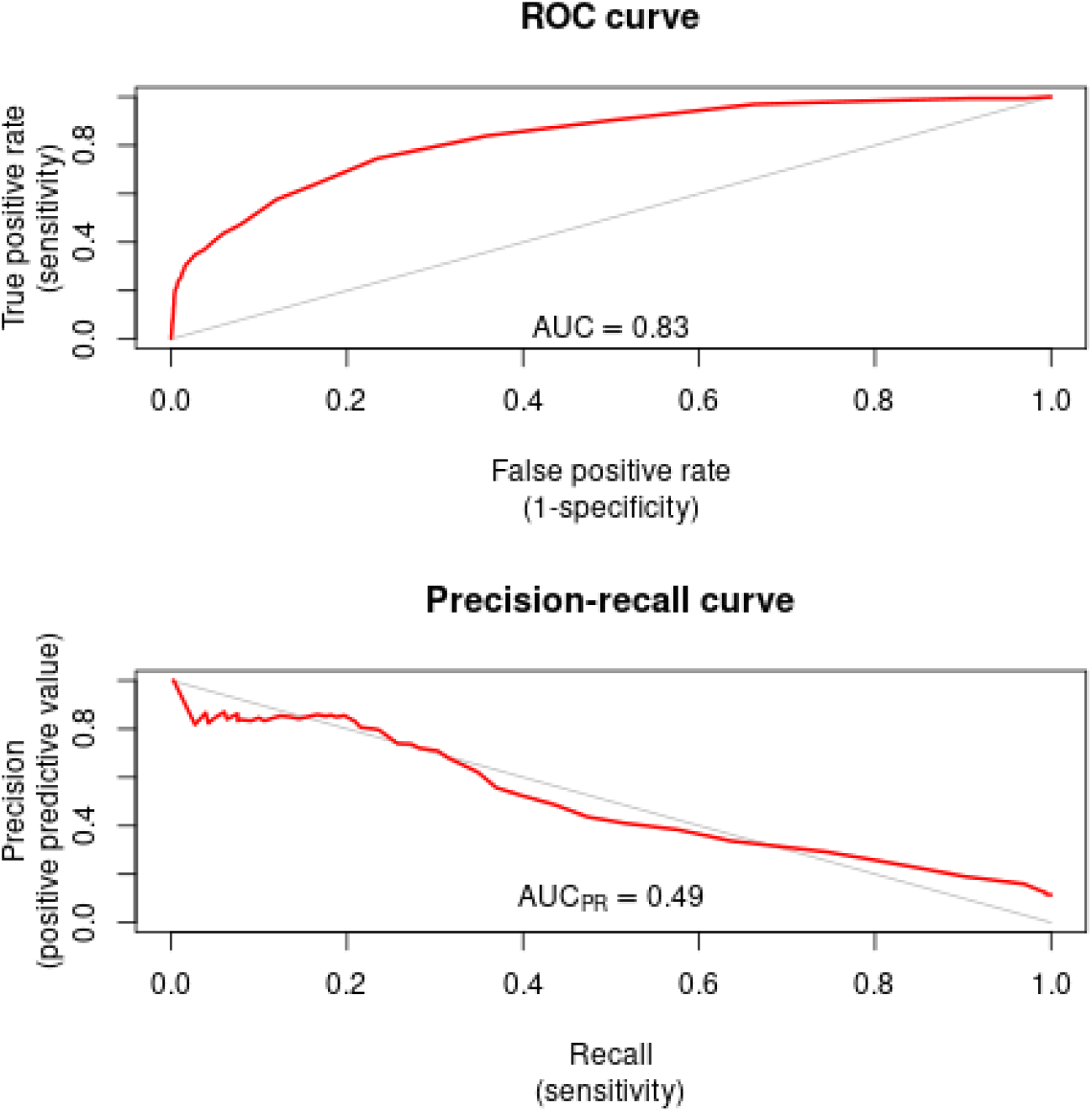
Synergy prediction using MLP. Evaluation of synergy prediction performance using ROC and PR Curves for a MLP.

**Table S1:**
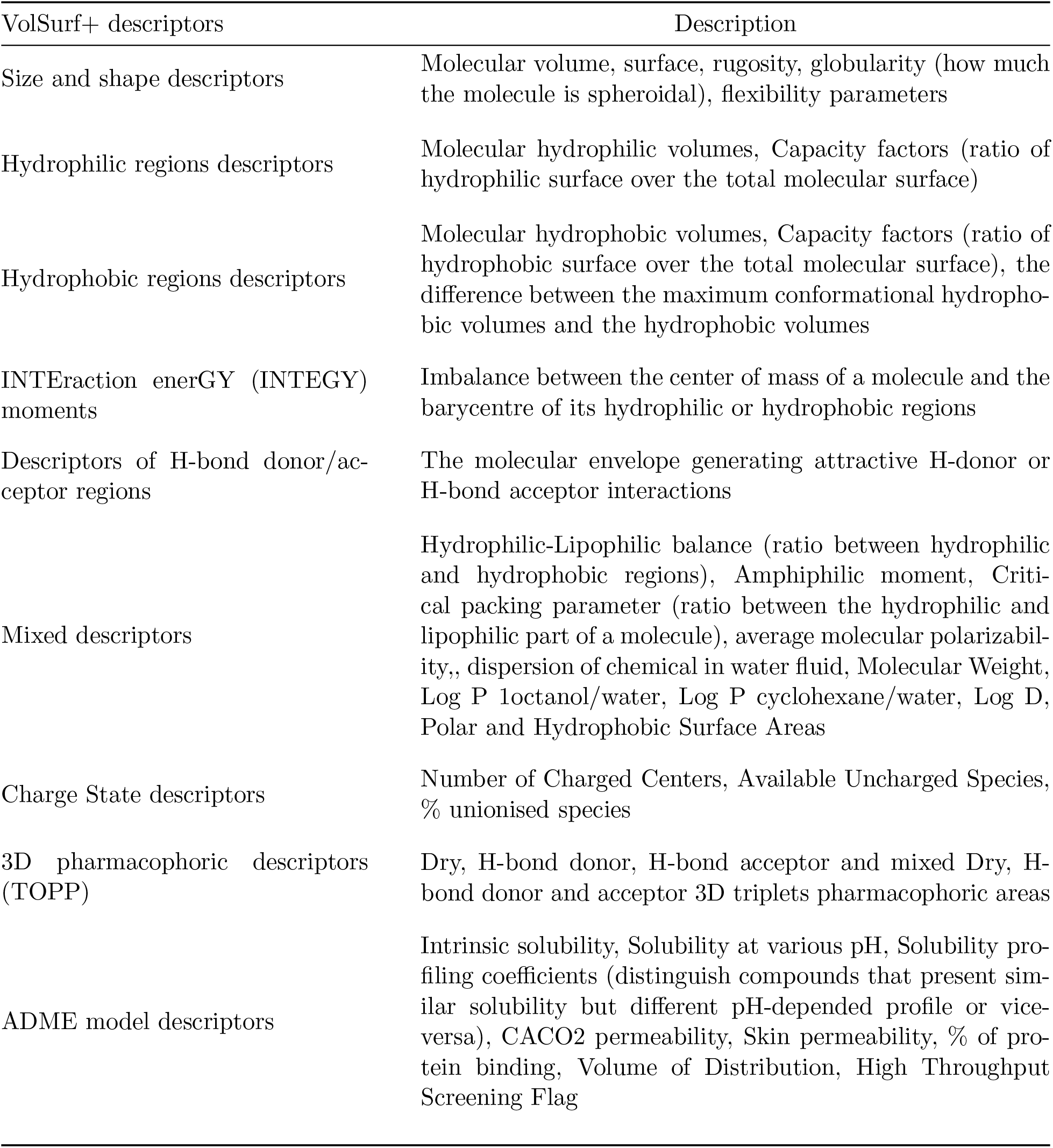
Illustration of VolSurf+ descriptors

**Table S2:** Comparison between MOViDA and DrugCell using: macroaverage MSE (MMSE) over all classes, lower classes (0-5), and higher classes (6-11), sampled Pearson correlation and sampled Spearman correlation. Each are average over k-fold cross-validation.

|  | MMSE | MMSE (0-5) | MMSE (6-11) | Pearson | Spearman |
| --- | --- | --- | --- | --- | --- |
| MOViDA | 0.025 | 0.032 | 0.020 | 0.89 | 0.89 |
| DrugCell | 0.035 | 0.060 | 0.018 | 0.86 | 0.88 |
| MOViDA leave-cell-lines-out | 0.040 | 0.059 | 0.026 | 0.82 | 0.83 |
| DrugCell leave-cell-lines-out | 0.054 | 0.100 | 0.021 | 0.81 | 0.82 |
| MOViDA leave-drugs-out | 0.098 | 0.160 | 0.055 | 0.51 | 0.52 |
| DrugCell leave-drugs-out | 0.128 | 0.195 | 0.080 | 0.40 | 0.50 |

## Notes

https://github.com/Luigi-Ferraro/MOViDA

https://doi.org/10.5281/zenodo.7406327

### Competing Interest Statement

The authors have declared no competing interest.

